# Primary visual cortex excitability is not atypical in acquired synaesthesia

**DOI:** 10.1101/814475

**Authors:** Laura Lungu, Ryan Stewart, David P. Luke, Devin B. Terhune

## Abstract

A wealth of data suggests that psychedelic drugs elicit spontaneous perceptual states that resemble synaesthesia although it is unclear whether these different forms of synaesthesia share overlapping neural mechanisms. Multiple studies have shown that developmental and trained synaesthesia is characterized by selective hyperexcitability in primary visual cortex and it has been proposed that cortical hyperexcitability may contribute to induced and acquired synaesthesia. This study tested the prediction that a case of acquired synaesthesia (LW) would display selectively elevated primary visual cortex excitability, as reflected in lower transcranial magnetic stimulation (TMS) phosphene thresholds, but no difference in motor thresholds, relative to controls. In contrast to this prediction, LW’s phosphene threshold was well within the threshold range of controls. These results suggest that acquired synaesthesia is not characterized by atypical visual cortex excitability.

---

Synaesthesia is a condition wherein stimulation of one modality automatically and consistently triggers activation of a secondary concurrent experience in a separate modality (Ward, 2013). Although it is typically viewed as a healthy neurodevelopmental condition with idiosyncratic inducer-concurrent associations that manifest in childhood, recent evidence suggests that synaesthesia-like experiences can be temporarily induced via consumption of recreational drugs, particularly serotonin receptor agonists (e.g., LSD, psilocybin) (Luke et al., 2013). Nevertheless, whether such experiences constitute a genuine form of synaesthesia remains controversial since drug-induced synaesthesias do not meet behavioural diagnostic criteria for developmental synaesthesia (Yanakieva, Luke, Jansari, & Terhune, 2019).

One interpretation of these phenomenological-behavioural discrepancies is that drug-induced synaesthesias do not meet such criteria because inducer-concurrent associations must undergo a process of consolidation before they achieve the level of automaticity and consistency characteristic of developmental synaesthesia (Terhune et al., 2016). We recently reported evidence in support of this hypothesis with a case of acquired synaesthesia (LW) following use of the recreational partial serotonin receptor agonist 2,5-dimethoxy-4-bromophenethylamine (2C-B; Yanakieva et al., 2019). LW has reported multiple forms of synaesthesia for over 9 years since ingesting 70-150mg of 2C-B, which is approximately 3-12 times the normal dosage (Johnson, Mathis, Shulgin, Hoffman, & Nichols, 1990). Using standardized measures, we corroborated that one or more of LW’s multiple forms of synaesthesia exhibited either consistency or automaticity, thereby meeting diagnostic criteria for this condition (Ward, 2013). Despite these results, it remains unclear whether acquired synaesthesia shares overlapping neural mechanisms with developmental synaesthesia.

It was recently proposed that drug-induced synaesthesia results from serotonin cascades triggering elevated cortical excitability in layer V pyramidal neurons, resulting in anomalous perceptual states that are mapped onto inducers, yielding synaesthetic experiences (Brogaard et al., 2013). This hypothesis is consistent with research showing selective hyperexcitability in primary visual cortex, as measured by transcranial magnetic stimulation (TMS) phosphene thresholds, but not motor (control) thresholds, in developmental synaesthesia (Terhune, Song, & Cohen Kadosh, 2015) and after synaesthesia training in controls(Rothen, Schwartzman, Bor, & Seth, 2018). If cortical hyperexcitability plays a role in induced synaesthesias, individuals with acquired synaesthesia will display selectively lower phosphene thresholds similar to developmental synaesthetes. The present study sought to test this prediction in LW using a double-blind design and standardized TMS protocols.

LW (31 years old, male) and 29 non-synaesthete controls (17 females, 12 males, *M*_Age_= 25.32, *SD* = 4.45) provided informed written consent to participate in this study in accordance with local ethical approval. LW did not significantly differ in age from controls, *t*=1.25, *p*=.11, *Z*_cc_=1.28 [0.83, 1.57] with a probability of occurrence in the general population of *p*_gp_=11% [7, 21]. None of the participants displayed contraindications for non-invasive brain stimulation and there were no adverse events.

Phosphene and motor threshold estimation followed established TMS protocols (Abrahamyan et al., 2011; Badran et al., 2019) (see **Supplementary Materials**). After identification of suitable stimulation sites, single-pulse TMS was applied to left primary motor cortex (motor thresholds) and midline primary visual cortex (phosphene thresholds) in counterbalanced order. For motor threshold estimation, participants rested their right hand on a table, with their forefinger and thumb touching and attended to the interosseous muscle. After stimulation to primary motor cortex with the coil positioned at a 45° postero-lateral angle, they reported whether they observed a visible twitch of the interosseous muscle during the stimulation. For phosphene threshold estimation, participants were first dark-adjusted and then sat with their eyes open at a 1m distance from a large, non-reflective black curtain (3 x 3 meters). After each stimulation with the TMS handle and coil in the vertical position, participants reported whether they had experienced a phosphene. Stimulation intensity varied on a trial-by-trial basis according to individual participants’ reports, as determined by a Bayesian adaptive staircase procedure with 30 trials run on each site in order to estimate 60% thresholds for both motor and phosphene thresholds (Abrahamyan et al., 2011).

As can be seen in **Figure 1**, in contrast to our central prediction, LW did not display a significantly lower TMS phosphene threshold (61) than controls, *M*=63.38, *SD*=9.49, *t*=0.25, *p*=.60, *Z*_cc_=0.25 [0.13, 0.58], with a probability of occurrence in the general population of *p*_gp_=60% [45, 72]. LW displayed a marginally higher motor threshold, 72 (corresponding to the highest threshold in controls), than controls, *M*=56.52, *SD*=7.51, *t*=2.03, *p*=.026, *Z*_cc_=2.06 [1.70, 2.41], *p*_gp_=3% [1, 5], and a marginally greater threshold difference (motor-phosphene), 11, relative to the controls, −6.86, *SD*=10.28, *t*=1.71, *p*=.049, *Z*cc=1.74 [1.37, 2.09] with a probability of occurrence of 5% [2, 9].

**Figure 1.**
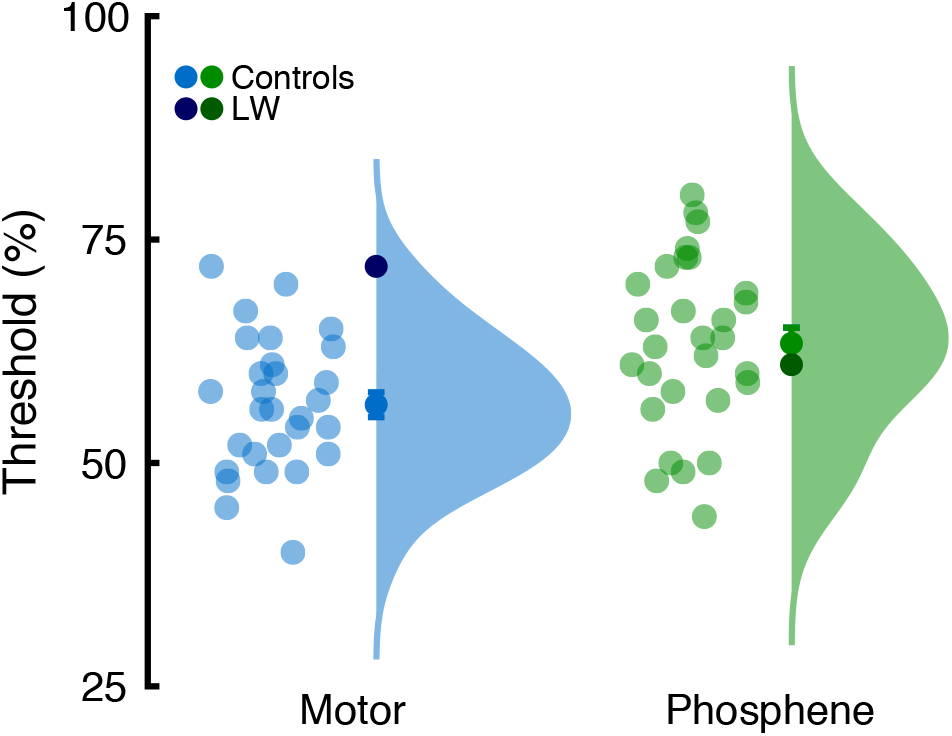
TMS motor and phosphene thresholds in controls and LW (acquired synaesthete). Error bars reflect standard error of the mean and marginal plots reflect kernel density plots.

We tested the prediction that a case of acquired synaesthesia would be characterized by selective hyperexcitability in primary visual cortex, as observed in developmental synaesthesia (Terhune et al., 2015) and induced synaesthesia (Rothen et al, 2018). In contrast with this prediction, we found that LW, an acquired synaesthete with previously-demonstrated inducer-concurrent automaticity and consistency (Yanakieva et al., 2019), displayed typical TMS phosphene thresholds that were within 25% of a *SD* of the mean phosphene threshold in controls. We did observe a tendency for LW to display marginally greater motor thresholds than controls, but this effect was unpredicted and should be interpreted with caution. These results suggest that LW does not exhibit primary visual cortex hyperexcitability and are at odds with the broader hypothesis of a role for cortical excitability in induced synaesthesia (Brogaard, 2013). One interpretation of these results is that atypical cortical hyperexcitability will only be observed in acquired synaesthesia further downstream in visual cortex, such as V4.

## Compliance with ethical standards

The authors report no conflicts of interests.

## SUPPLEMENTARY MATERIALS

### Methods

#### Participants

LW is a 31-year-old left-handed male and native English speaker with 3 years of higher education. LW has been experiencing multiple types of synaesthesia for the past 9 years after ingesting a 70-150mg of 2C-B (significantly more than the normal dose of 12-24 mg) at 22 years old. LW reported that his synaesthetic associations stabilized over time and his consistency scores on synaesthesia battery (Eagleman et al. 2007) met the criteria for week-colour and instrument-colour synaesthesia. Additionally, LW experiences visuospatially co-localized colours in response to face inducers (Yanakieva et al., 2019). 2C-B use was found to induce synaesthetic experiences in previous studies (Luke et al. 2012), but there have been no other reports of acquired synaesthesia in response to it to our knowledge. The control sample was comprised of 29 healthy adults without synaesthesia. No formal statistical power analysis was undertaken – the controls represented a convenience sample that were drawn from another as of yet unpublished study.

#### TMS Protocol

Participants wore a tight-fitting swimming hat made of lycra fabric, upon which circular stickers of 1 cm diameter were placed to indicate the optimal stimulation positions for the coil measuring motor and phosphene thresholds. The stimulation sites were determined measuring the scalp of each participant. TMS was applied using a DuoMag MP-Dual Paired Pulse TMS machine from Rogue Resolutions, with a DuoMag 70mm Butterfly Coil to administer the stimulation. Motor and phosphene threshold estimations were counterbalanced.

For motor thresholds, the left motor cortex was marked by moving 5 cm to the left and 2 cm anterior of the vertex. The observation of movement method was used in which participants reported whether they observed a twitch in the right hand interosseous muscle concurrent to the stimulation. This method was recently shown to be reliable compared to EMG methods (Badran et al., 2019). Prior to phosphene threshold estimation, participants were dark-adapted. For phosphene thresholds, the primary visual cortex was marked 2 cm above the inion (both measurements used in Terhune et al, 2015).

Once these locations were established, subsequent stimulations of and around the two sites were performed (moving no more than 1 cm at a time around the measured location) to identify sites that reliably elicited motor twitches or phosphenes, respectively. Threshold estimation was performed using a Bayesian adaptive staircase method (Abrahamyan et al 2011). After each stimulation, the participant was coded as displaying the response (twitch or phosphene) or not, and a new stimulation level was generated according to the adaptive staircase. A total 30 stimulation trials with varying stimulation intensities were applied, to estimate and record the 60% threshold.

#### Data analysis

LW and controls’ TMS thresholds and the difference between the two were contrasted using modified independent *t*-tests and effect size estimates (*z*_cc_) for case-control designs. In addition, for each effect, we estimated the probability of occurrence of the respective observation in the general population (*p*_gp_). We report bootstrap 95% confidence intervals for the latter two estimates (10,000 samples, bias-corrected and accelerated method) (Efron 1987). Raw data are presented in Table S1.

**Table S1.**
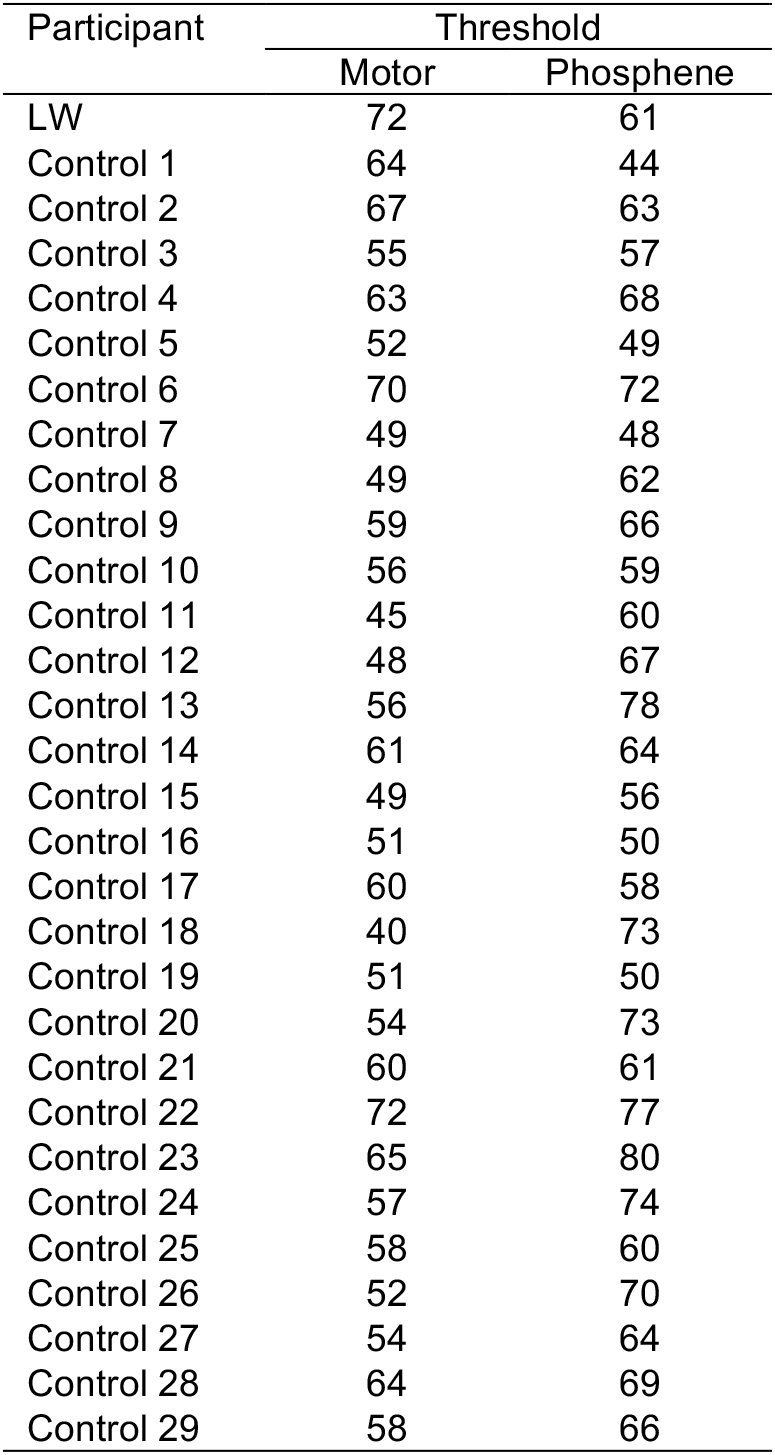
TMS thresholds in LW (acquired synaesthete) and controls.

